# The geographic distribution of reef and oceanic manta rays in the south-east Indian and south-west Pacific Oceans

**DOI:** 10.1101/727651

**Authors:** Amelia J. Armstrong, Asia O. Armstrong, Michael B. Bennett, Frazer McGregor, Kátya G. Abrantes, Adam Barnett, Anthony J. Richardson, Kathy A. Townsend, Christine L Dudgeon

## Abstract

The reef manta ray, *Mobula alfredi*, occurs in tropical and warm temperate coastal waters, and around islands and reefs in the Pacific and Indian Oceans. Published records that relate to the distribution of *M. alfredi* in the south-east Indian and south-west Pacific Oceans are largely restricted to locations where there is a focus on manta ray ecotourism, with little information from elsewhere. Even less is known about the circumglobally distributed oceanic manta ray, *Mobula birostris*, for which there are few published sighting records. We collated *n =* 11,703 sighting records from Australian waters and offshore territories for *M. alfredi* sourced from scientific image databases (*n* = 10,715), aerial surveys (*n* = 375) and online reports (*n* = 613). From collated records, we confirm that the species shows an uninterrupted distribution within Australian coastal waters north of 26°S on the west coast to 31°S on the east coast, with a southernmost record at 34°S. Confirmed locations for *M. birostris* encompass a latitudinal range of 10-40°S. Records from more southerly locations relate to warm-water events. Sightings of *M. birostris* were rare, but were confirmed at several geographically separate locations, probably reflecting its preference for offshore waters. The study clarifies the occurrence and range of each species within coastal waters of the south-east Indian and south-west Pacific Oceans, and highlights regions in northern Australia that are of specific interest for future research into possible movements of individuals between international marine jurisdictions.

## Introduction

The reef manta ray, *Mobula alfredi*, has a broad geographical distribution throughout much of the tropical and subtropical Indo-Pacific region, with the majority of records from relatively shallow waters associated with mainland coastlines, offshore islands, and reefs (Marshall, Compagno & Bennett, 2009; Couturier *et al.* 2012; Stewart *et al*., 2018) (Figure 1). However, the known distribution of *M. alfredi* is extremely patchy, with most records from dive ecotourism hotspots in Mozambique, South Africa, Maldives, Japan, Guam, the Red Sea, Philippines, New Caledonia, Indonesia and Australia (Figure 1). The distribution of *M. alfredi* could therefore be indicative of highly selective environmental preferences of this species or a consequence of sampling bias.

**Figure 1:**
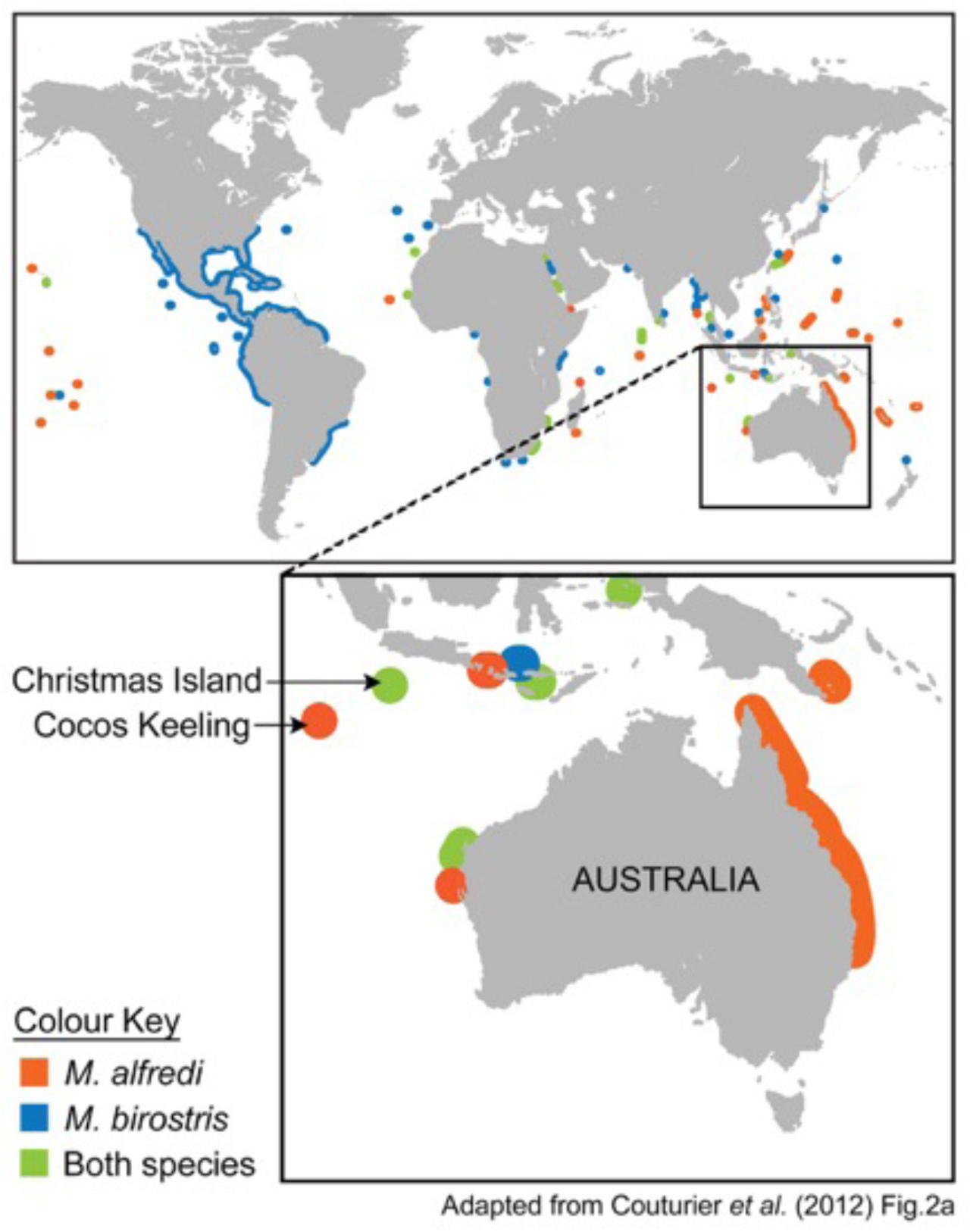
Previously published distribution for *M. alfredi* 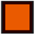 and *M. birostris* 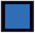 or both 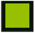, globally (main) and for the south-east Indian and south-west Pacific Oceans (inset). Note the species’ patchy distributions, including the discontinuity between the west and east coasts of Australia. Adapted from Couturier *et al.* (2012).

In the south-east Indian and south-west Pacific Oceans, the reef manta ray is thought to occur in shallow tropical and subtropical regions to a southern latitudinal extent of ∼30°S on the east coast of Australia and ∼26°S on the west coast (Last and Stevens, 2009; Couturier *et al.* 2012). Similar to the global distribution, the majority of these sighting reports are from subtropical locations where there are large seasonal aggregations of *M. alfredi* and regular ocean-based tourism activities. While *M. alfredi* has been reported widely along the eastern Australian coastline (Couturier *et al*., 2011; 2014), the occurrence of the species elsewhere remains mainly anecdotal.

Few records exist for *M. alfredi* from tropical Australian waters, likely due to a combination of relatively sparse human coastal populations, a large coastline with limited accessibility, regions of high turbidity, and a lack of in-water diving activities due to risks posed by the salt-water crocodile *Crocodylus porosus* and the box jellyfish *Chironex fleckeri*. However, given the adequately warm thermal environment and lack of barriers to movement along Australia’s northern coastal habitats, it is possible that *M. alfredi* has an uninterrupted distribution between the sub-tropical regions of both coastlines. Here we collate information from a variety of sources to examine the distribution of *M. alfredi* outside of known aggregation sites in Australian coastal waters. We also collated sighting information for *M. birostris*, which is based on relatively rare photographs of individuals at distinct locations (Figure 1; Couturier *et al.* 2015). Clarification of our understanding of the spatial distribution of these species provides a basis for future research into regional connectivity and conservation of sub-populations, and for evaluation of potential future changes in their geographical occurrence.

## Materials & Methods

We collated data on the locations of manta ray sightings from various sources, including image databases used for identification of individual manta rays, aerial surveys, reported online sightings, museum records, and the literature. All research complied with The University of Queensland’s Animal Ethics Committee approvals (SBS/319/14/ARC/EA/LEIER, SBS/342/17) and was conducted under the relevant marine park permits (Great Barrier Reef Marine Park Authority Permit G12/35136.1 and G16/37856.1).

### Manta database records

From birth, *Mobula alfredi* and *M. birostris* possess unique ventral patterning that has facilitated their inclusion in scientific image databases that record unique identifications and track records of individuals through time (Marshall & Pierce, 2012). Encounters with manta rays that have associated identification images are collated in two such image databases (one for the east and one for the west coast of Australia), both actively maintained by *Project Manta* (Couturier *et al*. 2011; McGregor *et al.* 2008). These databases contain images of manta rays encountered by researchers and citizen scientists. The latter contribution forms a large component of the east coast database and derives from social media engagement, or images and metadata submitted to *Project Manta* or other online observation portals such as Eye on the Reef (Dudgeon *et al.* in press; GBRMPA 2018). Electronic submission is typically an image of the ventral body surface of a manta ray, along with sighting location, date of capture, and behavioural observations. Each image is scrutinized and matched to the ventral body surface pattern specific to an individual manta ray if that individual has been photographed previously. A novel body surface pattern indicates a previously unrecorded individual, and is assigned a new, unique, identification code.

### Aerial surveys

Between November 2017 and February 2018, we conducted aerial surveys targeting manta rays on near-shore reefs around Cairns (16.92° S, 145.78° E) and Port Douglas (26.48° S /145.46° E) Queensland, using GLS Aviation (https://gslaviation.com.au/cairns/). Sightings including the number of individuals and GPS location were reported by pilots during scheduled flights covering two scenic tourism routes (https://gslaviation.com.au/cairns/reefhopper).

We also collated data on manta ray presence from population monitoring surveys that targeted marine mammals such as dugongs *Dugong dugon* and various dolphin species. Aerial surveys with sightings of manta rays cover Ningaloo Reef and Shark Bay in Western Australia (Preen *et al.* 1997, Hodgson 2007) and the coastline of the Northern Territory (Groom *et al.* 2015, Palmer 2015, Groom *et al.* 2017).

### Online sighting records

Additional manta ray location records were found through extensive online searches (Google, www.google.com; YouTube, www.youtube.com). We used species keywords (e.g. manta, reef manta) in combination with specific locations (e.g. Rowley Shoals, Dampier, Broome, Darwin, Weipa, Groote Eylandt, among others), regions (e.g. Northern Territory, far North Queensland) and activity (e.g. snorkelling, fishing, fly fishing), to identify potential sighting records of *M. alfredi* around Australia. The online search yielded results across a variety of sources including the scientific literature, business and personal blogs, fishing forums, local news reports, videos and online image galleries (e.g. Eye on the Reef, GBRMPA; Reef tourism operator image galleries). As there could be misidentifications (e.g. different species of *Mobula* and eagle rays might be reported as *M. alfredi*), sighting reports were classified as those with or without supporting imagery. If imagery to support a manta ray sighting was lacking, reports were only included if accompanying descriptions (e.g. swimming at the surface with mouth open, circling in dense patches of plankton; or body size estimates, e.g. 2 – 4 m disc width) matched known *M. alfredi* behaviours and/or morphology.

## Results

In total, we collated 11,703 sighting records for *M. alfredi* in Australian waters from the various data sources. The species showed a near-continuous distribution across northern Australian coastal waters from Shark Bay in Western Australia to the Solitary Islands Marine Park in New South Wales (Table 1; Figure 2). We also found 29 sighting records for the oceanic manta ray *M. birostris* (Figure 2).

**Table 1:** Collated sightings for *Mobula alfredi and M. birostris* in Australian waters summarised by source.

| **Sighting Record Source** | *Mobula alfredi* | *Mobula birostris* |
| --- | --- | --- |
| Manta ray identification database | **10,715** | **16** |
| *Project Manta* East coast database | **6480** | **2** |
| *Researcher* | 1826 |  |
| *Citizen Scientist* | 4200 |  |
| *Researcher & Citizen Scientist* | 454 |  |
| *Project Manta* West coast database | **4235** | **14** |
| *Researcher* | 2127 |  |
| *Citizen Scientist* | 2108 |  |
| Aerial Observations | **375** | **2** |
| *Marine Mammal Survey* | 255 | 2 |
| *Manta Survey* | 120 |  |
| Online sighting reports | **613** | **7** |
| *No supporting media* | 492 | 2 |
| *Supporting media (e.g. image, footage)* | 121 | 5 |
| **Museum records** | **0** | **1** |
| **Total** | **11,703** | **26** |

**Figure 2:**
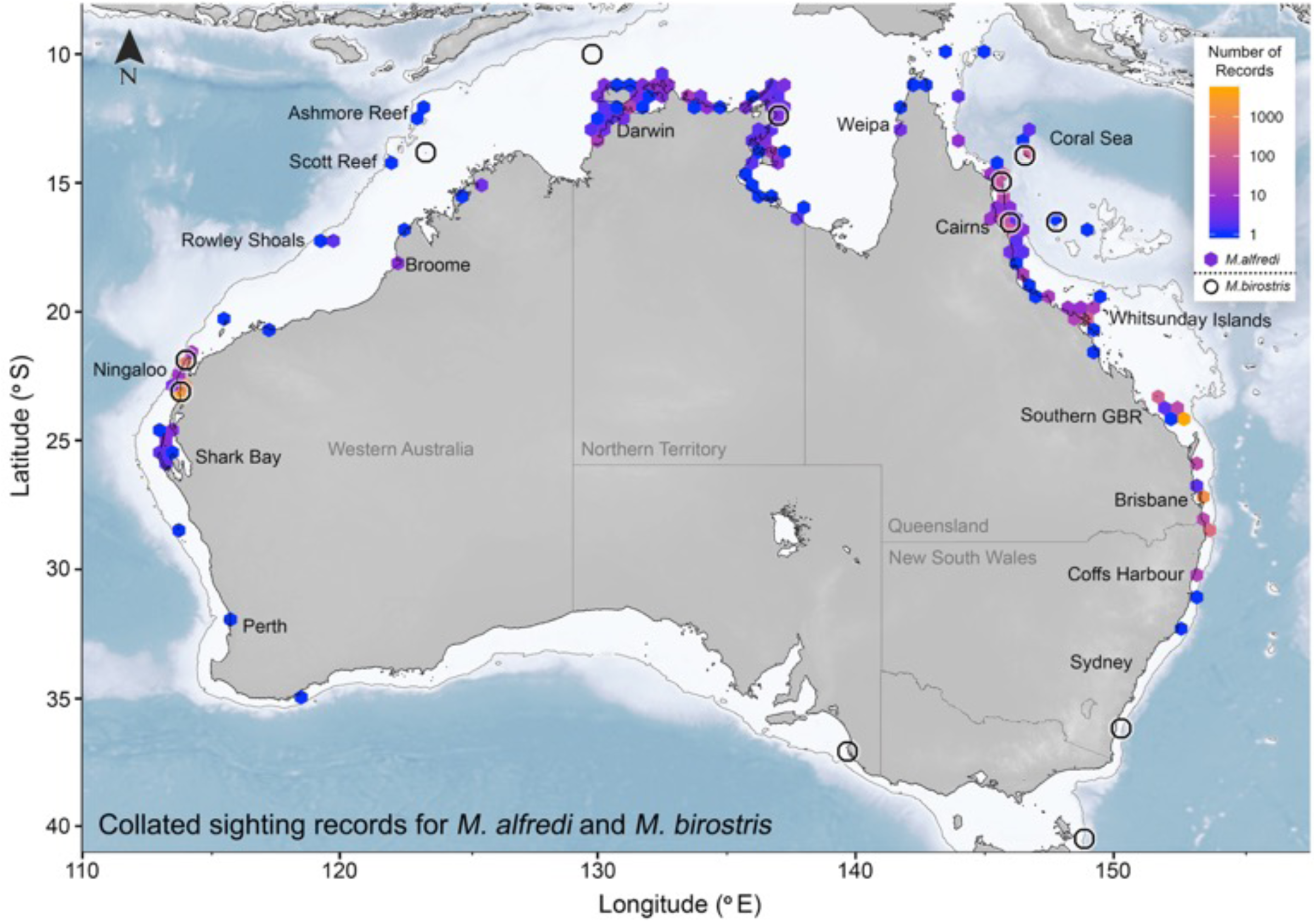
Collated sighting records for *Mobula alfredi* and *M. birostris* in Australia, with data sourced from scientific literature, image databases, aerial surveys, museum records, and online reports. *Mobula alfredi* sightings are aggregated across a 0.5 ° gridded area and are represented by hexagonal cells, where colour is indicative of the sighting count (from 1 to >1000) per cell. Sightings of *M. birostris* are represented by open black circles and are indicative of location only (not count, due to the limited number of observations). Note that data from Cocos Keeling Island and Christmas Island are not shown. The unbroken grey line off the coast of Australia represents the 500 m isobath.

### Manta database records

Over 90% of the collated sighting records were sourced from the two manta identification databases maintained by the *Project Manta* research group (*n =* 10,715) (Supplementary). The east coast database contained ∼55% (*n =* 6,480) of the sightings, with *M. alfredi* observed between 13- 31°S. However, 91% of those reports were from just two locations (Lady Elliot Island = 71%; North Stradbroke Island = 20%).

Along Australia’s west coast, photographic observations of manta rays are largely restricted to the Ningaloo coastline where on-water recreation and aquatic ecotourism activities occur year-round (Smallwood *et al.* 2011, Venables *et al.* 2016). Outside of the Ningaloo Reef region, vast tracts of coastline are uninhabited and inaccessible for tourism activity. The west coast database contained 36% (*n* = 4,235) of Australia’s sighting records, with all but three sightings from the Ningaloo coastline, between 21.6°S to 23.5°S. Citizen science submissions constituted 64% (*n* = 4200) and 50% (*n* = 2108) of east and west coast database entries respectively. There were 16 database records for *M. birostris*, 14 of which were from the Ningaloo Reef region, with the remaining two from the Great Barrier Reef and Coral Sea.

### Aerial records

Although aerial records represented just 3% (*n* = 375) of the total Australian manta ray sighting records, they filled in much of the previously unsampled area in northern Australia. Mobulids, most likely *M. alfredi*, were seen along the Northern Territory coastline, with additional localized coverage and reports from Western Australia and Queensland (Supplementary). For aerial surveys targeting manta rays, 95 *M. alfredi* were observed on 48 scenic reef flights by GSL aviation and 25 *M. alfredi* from 3 research specific flights covering near-shore reefs of Cairns and Port Douglas. Northern Territory marine megafauna monitoring surveys (between 2014-2016) contributed 211 sighting records. Two of these records from the Northern Territory were identified by aerial observers as likely sightings of *M. birostris*. Similar aerial surveys covering the Ningaloo and Shark Bay regions in Western Australia (1989, 1994, 2007) contributed 44 records.

### Online records

Online sighting reports (5%; *n =* 613) provided evidence of manta ray presence in some areas that were not encompassed by the identification databases or aerial surveys (Supplementary). While 80% of those records lacked supporting media, the vast majority of observations were from within the Great Barrier Reef Marine Park where experienced tourism operators have a good knowledge of local species and *M. alfredi* is found; species identity was assumed to be correct given these circumstances. The remaining 20% of records with accompanying images that confirmed the identification of individuals as *M. alfredi* were generally near ‘remote’ towns (e.g. Weipa, Queensland), where there are on-water activities such as fishing. Seven records of *M. birostris* were found, five of which had supporting media to confirm the identification.

The most southerly records of *M. alfredi* on the west coast were from Coogee Beach, Perth (32.1°S) – with supporting image evidence – and Cheynes Beach, Albany (34.9°S) – without image-based confirmation of identity. On the east coast, the most southerly confirmed records are from South West Rocks, NSW (30.9°S). There were a few possible sightings of *M. alfredi* further to the south, but these, and a single record from Sydney Harbour in 1868, are excluded due to uncertainties about the species involved. Sightings were also recorded from Australia’s offshore remote territories in the Indian Ocean where presence of *M. alfredi* (Cocos Keeling Island) and *M. birostris* (Christmas Island) are confirmed.

## Discussion

By combining multiple data types, we produced the most comprehensive distribution map for *M. alfredi* within the south-east Indian and south-west Pacific Oceans, providing a significant update on the published species distribution. Rather than having highly selective environmental preferences that cause a patchy distribution, we consider that *M. alfredi* has a near-continuous distribution throughout Australia’s tropical and subtropical coastal waters southwest to about Shark Bay, Western Australia, and southeast to the Solitary Islands, New South Wales. We also confirm that *M. birostris* is found on all coasts of Australia, including in temperate waters of Tasmania (Couturier *et al.* 2015), but is much less common than *M. alfredi*.

The southerly range limit for *M. alfredi* is similar on both east and west Australian coasts. Australia is unique in being the only continent with warm poleward-flowing currents on both coasts: the East Australian Current in the east (Ridgway & Godfrey 1997) and the Leeuwin Current in the west (Godfrey & Ridgway 1985). These currents set the southern range of various mobile species (Couturier *et al.* 2011, Dudgeon *et al.* 2013, Payne *et al.* 2018). Many tropical marine fauna reach the southern limit of their range, and many temperate fauna reach their northern limit, at latitudes of ∼25°S to ∼30°S on Australia’s east and west coastlines (Gomon *et al.* 2008, Last & Stevens, 2009, Blair *et al.* 2014).

Given the few records of *M. alfredi* at locations further south than Coffs Harbour (∼25°S) or Shark Bay (∼30°S) on the east and west Australian coasts, respectively, it is likely that sightings south of these regions are unusual forays into normally cold-temperate environments and not part of their normal distribution. For example, manta rays (probably *M. alfredi*) reported as far south as Cheynes Beach (34.9°S, Albany, WA) coincided with an exceptional ‘marine heatwave’, when sea surface temperatures peaked at up to 5°C warmer than normal (Feng *et al.* 2013), and temporary range extensions of many marine fishes were recorded (Pearce & Feng 2013).

This study identified locations within Australia’s northern waters where *M. alfredi* have not been reported previously, including the coastline of the Northern Territory, the northern coasts of Western Australia and Queensland. Both reports and confirmed photographic identifications at remote Australian offshore territories of Christmas Island (*M. birostris*) and Cocos Keeling islands (*M. alfredi*), located ∼1500 and ∼2000 km offshore respectively. Our findings of manta ray distribution at the Cocos Keeling Islands and Christmas Island support those of Kashiwagi *et al.* (2011), with evidence for only *M. alfredi* at Cocos Keeling Islands and *M. birostris* at Christmas Island. These results differ to the distribution presented by Couturier *et al.* (2012) for Christmas Island, where both species were reportedly present.

The near-continuous distribution for *M. alfredi* raises questions about regional population structure. It is well-established that *M. alfredi* exhibits strong migratory behaviour, with movements of up to 500 km not uncommon (Couturier *et al.* 2011, Germanov and Marshall 2014, Jaine *et al.* 2014) and recently movements of at least 1150 km have been demonstrated (Armstrong et al. 2019). Photo-identification and tagging studies show that individual rays have an affinity for particular sites and regions (Braun *et al.* 2015, Couturier et al 2014, Kessel *et al.* 2017, Marshall *et al.* 2011). However, most studies have focused on populations separated by large distances and/or by regions of deep water (Deakos *et al.* 2011). The shallow continental shelf of northern Australia may faciliate broad-scale connectivity along the expansive coastline, given an apparent absence of barriers to movement. Whether *M. alfredi* move between Australia and its northern neighbours is curently unknown. Our study presents a single record from Ashmore reef, located ∼135 km south of Indonesia’s southernmost islands, Rote and Pulau Ndana. Distances between Ashmore Reef, the southernmost Indonesian islands, and the Australian mainland, fall within the known range of reef manta movement capacity (<500 km). Further collaborative work would be necessary to determine the potential for exchange of Australian genetic stock with that of Indonesian manta rays.

This study provides evidence that *M. alfredi* is distributed across nearly two-thirds of Australia’s coastline and adjacent islands and reefs. The species’ distribution is predominantly restricted to warm waters north of ∼30°S. Unusual events in which warm water extends further south than normal appear to be accompanied by temporary southerly range extensions. Although records are scarce, we provide the first map of *M. birostris* occurrence in Australian waters and confirm that this species is found on all coasts. Due to the proximity of some sightings to international marine jurisdictions and the species’ capacity to traverse the separating distances, future collaborative work is necessary to determine whether international movements occur.

## Acknowledgements

The authors wish to acknowledge the Northern Territory Department of Environment and Natural Resources for providing access to the dugong aerial survey data; GSL Aviation, Cairns, for assisting with the targeted aerial survey for manta rays; the Great Barrier Reef Marine Park Authority for access to manta ray data mined from the Eye on the Reef database, and finally we would like to thank the hundreds of citizen scientists from across Australia who contribute data to the *Project Manta* image databases.

## Funding information

This research was funded by the Australian Research Council Linkage Grant LP110100712. AJA & AOA were supported by a University of Queensland Research Scholarship.

## Contributions

The study was conceived by A.J.A, A.O.A and C.LD. All authors participated in data collection. The figures were produced by A.J.A and all authors contributed to manuscript creation. All authors have seen and approved the final manuscript.

## Significance Statement

Published records relating manta ray (*Mobula alfredi & M. birostris*) distribution in the south-east Indian and south-west Pacific Oceans are largely restricted to manta ray focused ecotourism localities, with little information from elsewhere. This study clarifies the occurrence and range of manta rays within coastal waters of the south-east Indian and south-west Pacific Oceans, demonstrating the value of citizen science and non-traditional data sources (social media and online videos) for understanding species distributions.

