## Supplemental Figures for "The geographic distribution of reef and oceanic manta rays in the south-east Indian and south-west Pacific Oceans"

*Corresponding author: amelia.armstrong[at]uqconnect.edu.au

**Supplementary Figures A-C:**

Sighting records for manta rays in Australia, mapped by data source. A = from scientific image databases, B = aerial surveys, and C = online reports. Sightings are aggregated across a 0.5° gridded area and represented by hexagonal cell, where colour is indicative of the sighting count (from 1 to >1000) per cell. Note that the colour scale representing the count of each hexagonal point is not equivalent across data sources and has been generated to best visualize sighting variance. Note that data from Cocos Keeling Island and Christmas Island are not shown. The unbroken grey line off the coast of Australia represents the 500 m isobath.

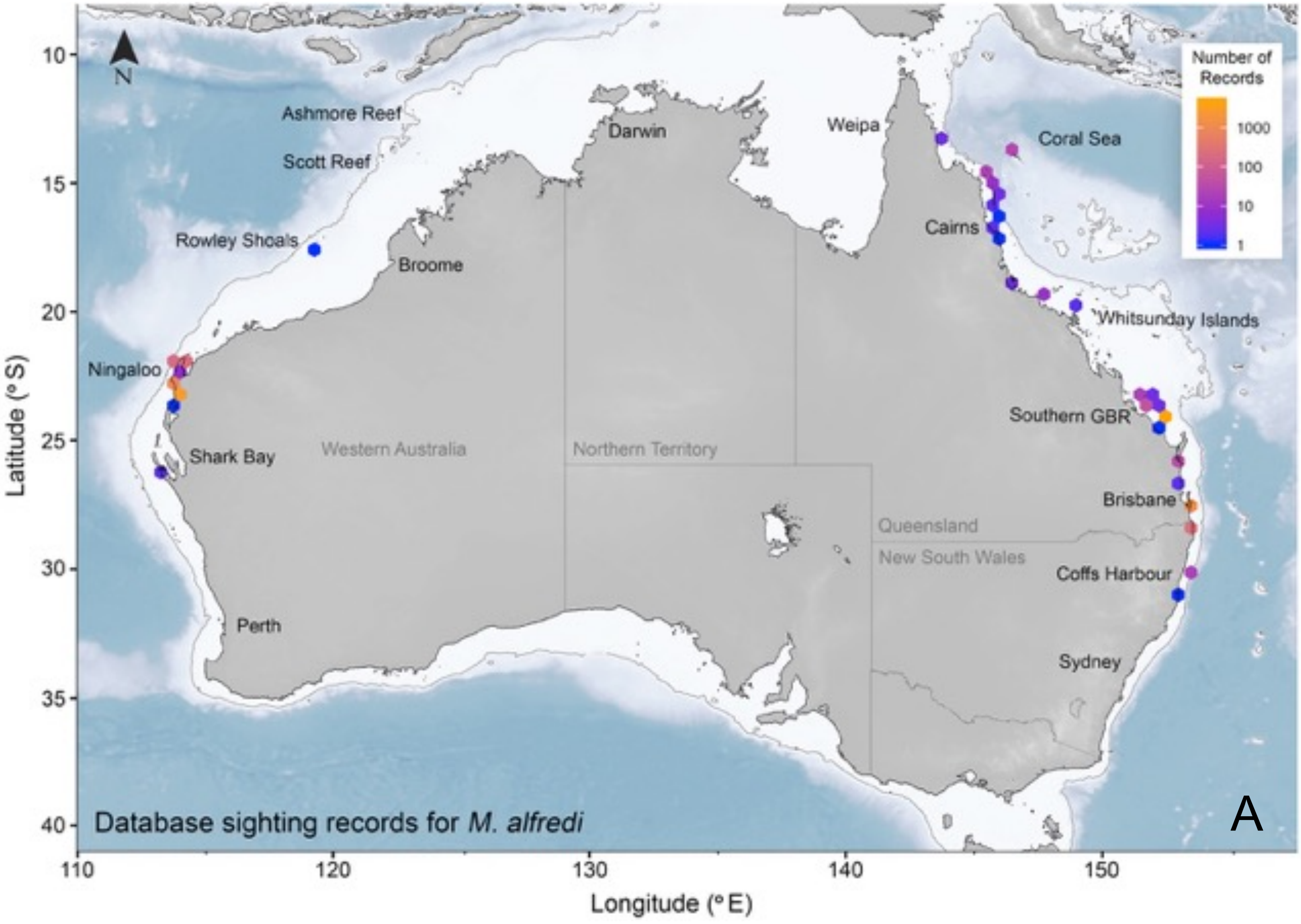

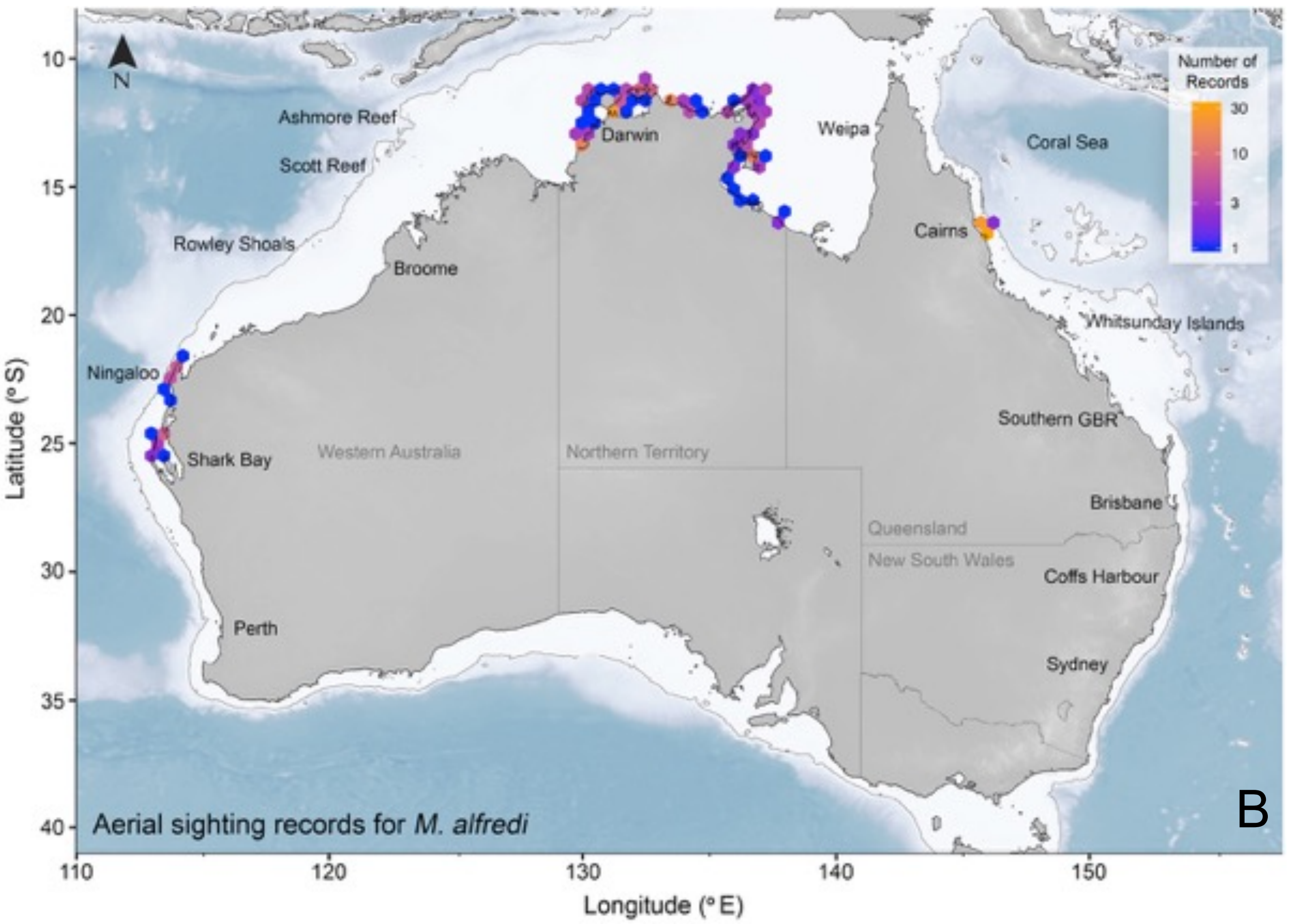
Number of Records
30
10
3
1
Ashmore Reef
Scott Reef
Rowley Shoals
Broome
Darwin
Weipa
Coral Sea
Cairns
Whitsunday Islands
Ningaloo
Shark Bay
Western Australia
Northern Territory
Southern GBR
Brisbane
Queensland
New South Wales
Coffs Harbour
Perth
Sydney
Aerial sighting records for M. alfredi
B
Latitude (° S)
Longitude (° E)
10
15
20
25
30
35
40
110
120
130
140
150

15

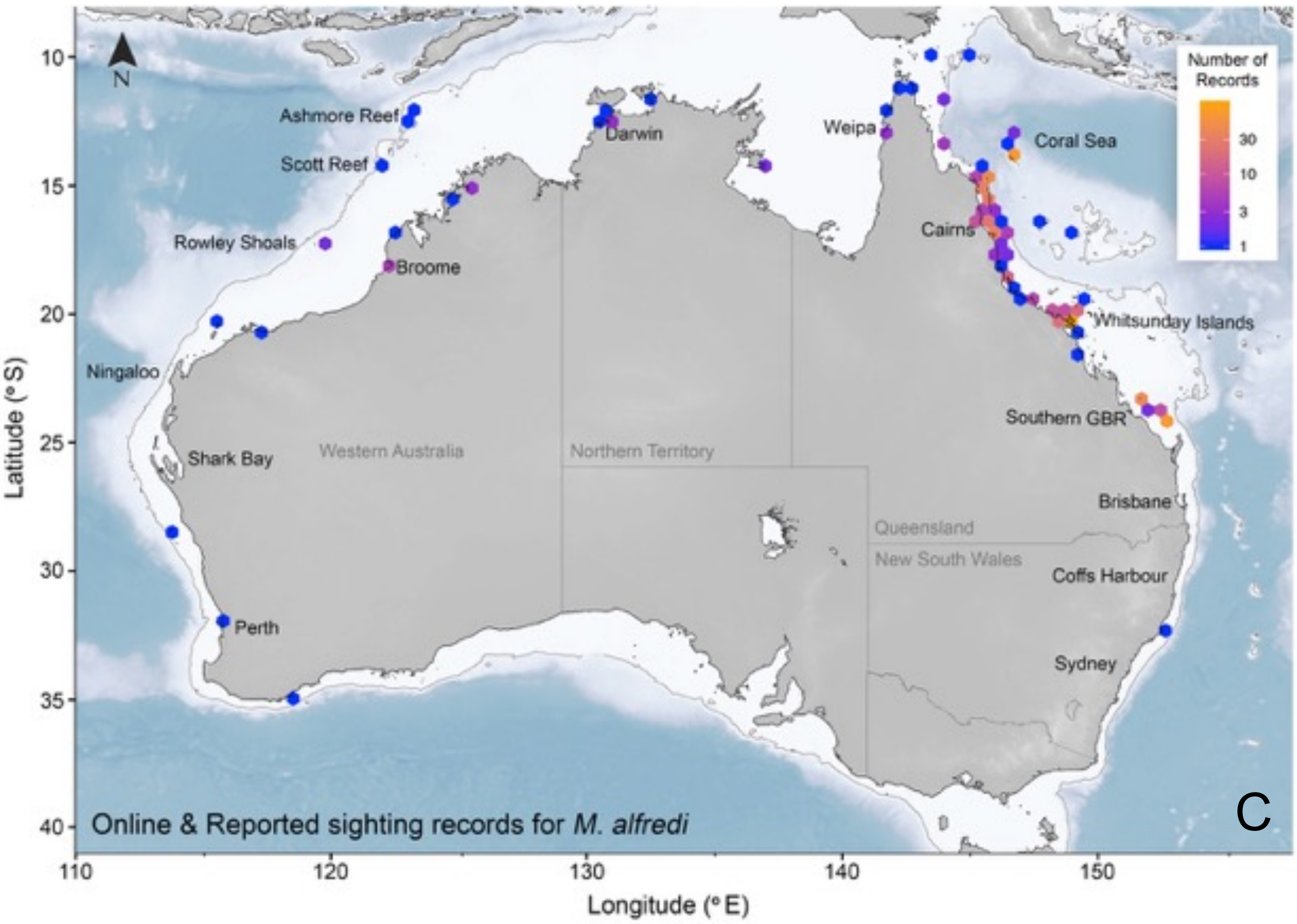
Number of Records
30
10
3
1
Ashmore Reef
Scott Reef
Rowley Shoals
Broome
Darwin
Weipa
Coral Sea
Cairns
Whitsunday Islands
Ningaloo
Shark Bay
Western Australia
Northern Territory
Southern GBR
Brisbane
Queensland
New South Wales
Coffs Harbour
Perth
Sydney
Online & Reported sighting records for M. alfredi
C
Latitude (° S)
Longitude (° E)
10
15
20
25
30
35
40
110
120
130
140
150
